# KOnezumi-AID: A Software to Automate the Design of Multiplex Knockouts Using Target-AID

**DOI:** 10.1101/2024.10.29.620976

**Authors:** Taito Taki, Kento Morimoto, Seiya Mizuno, Akihiro Kuno

## Abstract

With the groundbreaking advancements in genome editing technologies, particularly CRISPR-Cas9, creating knockout mutants has become highly efficient. However, the CRISPR-Cas9 system introduces DNA double-strand breaks, which can increase the risk of chromosomal rearrangements, presenting a major obstacle when attempting to knockout multiple genes simultaneously. Base editing systems like Target-AID provide a safer alternative by enabling precise base modifications without requiring DNA double-strand breaks, making them a promising solution for addressing this challenge. Nevertheless, the absence of adequate tools to support Target-AID-based gene knockouts highlights the need for a comprehensive system to design guide RNA (gRNA) for the simultaneous knockout of multiple genes. Here, we present KOnezumi-AID, a command-line tool designed to facilitate gRNA design for Target-AID-mediated genome editing. KOnezumi-AID supports gene knockout strategies by inducing premature termination codons or promoting exon skipping. It generates experiment-ready gRNA designs for both mouse and human genomes. Additionally, KOnezumi-AID offers batch processing capabilities, enabling rapid and precise gRNA design for large-scale genome editing projects such as CRISPR screening. In summary, KOnezumi-AID provides an efficient and user-friendly tool for gRNA design, streamlining genome editing workflows and advancing gene knockout research.

## 1. Introduction

Since the establishment of clustered regularly interspaced palindromic repeat (CRISPR)-CRISPR-associated protein 9 (Cas9) as a genome editing technology, the speed of generating genetically modified mice has increased markedly [1,2]. According to the International Mouse Phenotyping Consortium, as of 2021, knockout (KO) mice have been produced for approximately 11,241 protein-coding genes [3], which accounts for nearly half of all mouse genes, and this number continues to grow. It is predicted that single-gene KO analyses will be largely completed in the near future, and the simultaneous creation and analysis of multiplex KO mutants will become a critical challenge [4,5].

However, KOs with CRISPR-Cas9 system involve the risk of inducing large-scale chromosomal deletions spanning several thousand base pairs, triggered by DNA double-strand breaks (DSBs) [6]. This risk increases significantly when targeting multiple genes simultaneously [7]. To address this issue, we focused on base editing, particularly Target-AID, a synthetic complex of Cas9 derived from *Streptococcus pyogenes* (SpCas9) fused to an activation-induced cytidine deaminase (AID). This enables gene editing without the need for DSBs. Target-AID is a type of base editor that can convert cytosine to thymine (C-to-T), introducing point mutations specifically at 17-19 nucleotides (nt) upstream of the protospacer adjacent motif (PAM) sequence [8]. Compared to other base editors, Target-AID has a narrower editing window [9], offering enhanced precision and reducing the risk of unintended mutations. Additionally, while other base editors have been associated with off-target effects on RNA, Target-AID poses a lower risk [10], making it a safer option for the simultaneous editing of multiple genes.

Strategies for achieving KO through the use of base editors include inducing premature termination codons (PTCs) to truncate mRNA [11,12], or introducing frameshift mutations via splice-site disruption [13,14]. However, the design of guide RNAs (gRNAs) for these strategies has required manual creation for each gene, which presents a significant obstacle to the design of experiments. Therefore, this study aims to develop ’KOnezumi-AID’, a tool that allows the automatic design of experiment-ready gRNAs for Target-AID simply by inputting the target genes to be knocked out. By requiring the gene symbols as input, this tool facilitates the rapid design of genome editing strategies, even for the KOs of multiple genes.

## 2. Results

### 2.1. Implementation of KOnezumi-AID

We implemented a command-line tool, KOnezumi-AID, to design optimal gRNAs for inducing KO of the protein-coding gene in mice using SpCas9 based Target-AID. We initially focused on mouse because it enables the rapid generation of KO mutants and has abundant publicly accessible genomic data [15, 16]. KOnezumi-AID requires the user to provide a refFlat file and a reference sequence as inputs for preprocessing. To extract gRNA candidates for protein-coding exons, KOnezumi-AID filters the refFlat file by removing non-protein-coding transcripts and duplicated transcripts. In addition, KOnezumi-AID targets both autosomes and sex chromosomes, excluding alternative assemblies (see details in Materials and Methods). Next, the sequences of the transcripts in the transcriptional direction are extracted according to the genomic annotations of the filtered refFlat file and saved as refseq_to_transcribed_regions.pkl. Moreover, to standardize subsequent computational processes, the genomic coordinates in each transcript are converted into relative coordinates starting from the transcribed regions using bedtools [17], and they are cached as unique_coding_refflat.pkl (Figure 1A).

**Figure 1.**
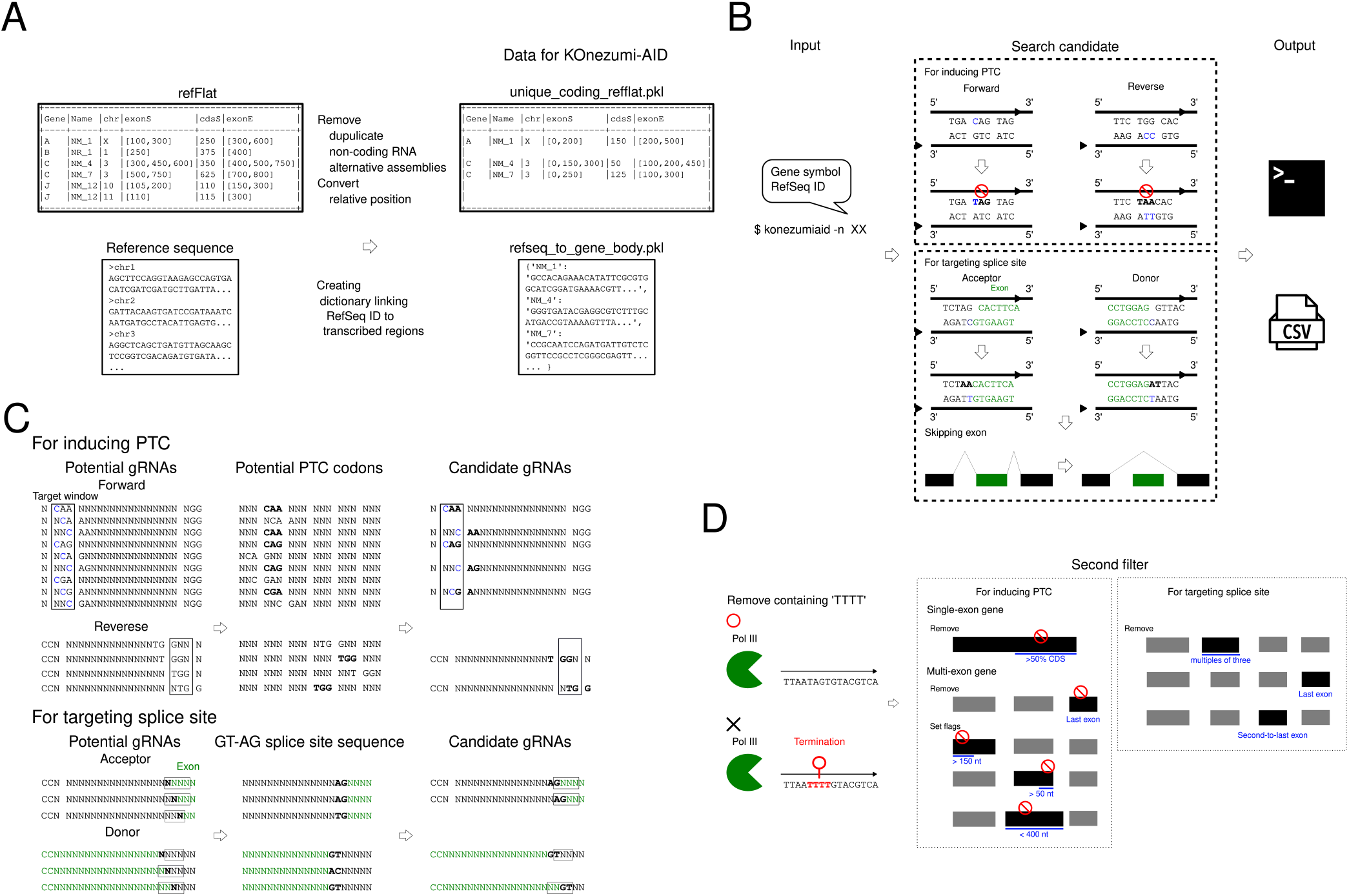
Overview of KOnezumi-AID: (A) Preprocessing in KOnezumi-AID. RefFlat files and reference sequences are processed into a format suitable for KOnezumi-AID searches. (B) The main search process of KOnezumi-AID. The blue highlights indicate the editable base. The bold high-lights and red stop marks indicate the PTCs. Exons are marked in green. (C) Search process in KOnezumi-AID. This schema represents the search process in KOnezumi-AID. The black box indicates the target window. The blue indicates editable bases. The bold highlights represent the regions where codons can be modified to introduce PTCs or target the GT–AG splice site consensus sequence. Exons are marked in green. (D) Filtering process in KOnezumi-AID. The red stop marks indicate PTCs, and the blue highlights represent key filtering checkpoints.

After preprocessing, KOnezumi-AID accepts a gene symbol or RefSeq ID as input from the user and identifies optimal gRNA candidates for knocking out the target gene. The gRNA design strategy of KOnezumi-AID focuses on inducing a PTC or causing a frameshift mutation by disrupting splice acceptor or donor consensus nucleotides [11-14]. The experiment-ready gRNAs are printed to the standard output and saved as a comma-separated values (CSV) files in the directory konezumiaid_data/output (Figure 1B).

Regarding the details of gRNA design, the potential gRNA candidates are first comprehensively listed. Considering that the target window for Target-AID is the cytosine located at +17 to +19 nt from the SpCas9-dependent PAM (NGG) [7], sequences that contain both a targetable cytosine and a PAM at exon or splice sites are extracted. To induce PTC, the target cytosine must be within the codons CAG, CGA, or CAA. For splice sites, ensure that a PAM is located +17 to +19 nt. For the purpose of inducing PTC, codons are calculated using the relative position of the ORF, and candidate gRNAs that result in a stop codon through C-to-T conversion are identified. When targeting splice sites, surrounding sequences at the exon start and end points are retrieved, and gRNAs targeting nucleotide conversion of the splice donor or acceptor are identified (Figure 1C).

After listing all candidate gRNAs, KOnezumi-AID filters them to identify experiment-ready gRNAs. In the first filtration step, gRNAs containing TTTT in the target sequence are excluded, as the use of pol III promoters for gRNA expression is a common vector-based approach, and the TTTT sequence triggers pol III termination [18]. Next, we established rules to extract gRNAs that are likely to induce a KO. For single-exon genes, we selected gRNAs that induce PTCs within the first 50% of the coding sequence (CDS) as candidates [15]. For multi-exon genes, we prioritized triggering nonsense-mediated mRNA decay (NMD), which is an mRNA quality control mechanism that degrades transcripts with abnormal stop codons, via PTC induction [19]. The gRNAs that induce PTCs in the last exon were excluded, as they do not lead to NMD [20,21]. Next, flags were assigned based on the following criteria [22]: the start 150 nt rule, which suggests that NMD is less likely if the PTC occurs within 150 nt of the start codon [22,23]; the last 50 nt rule, indicating that NMD is less likely if the PTC is within 50 nt of the last exon junction complex (EJC) [24]; and the long exon rule, where exons longer than 400 nt tend to suppress NMD [25]. For gRNAs targeting splice sites, those affecting exons whose lengths are multiples of three were excluded, as no frameshift would occur. Furthermore, gRNAs targeting the last exon were excluded to avoid potential splicing into other genes. Finally, gRNAs targeting the second-to-last exon were excluded, as PTC induction through frameshifts in this region would not trigger NMD (Figure 1D).

The output of KOnezumi-AID includes the gRNA sequence, PAM sequence, a link to CRISPRdirect [26], and the targeted amino acid for gRNAs designed to induce PTC. If the gene has multiple isoforms, gRNAs that can target all isoforms are output (Figure 2A). For input genes with a single transcript or those specified by RefSeq ID, the amino acid number of the target site is displayed. Additionally, if the gene is a multi-exon gene, a ’Recommended’ column is included, marked as True when all four criteria for NMD susceptibility are met, indicating a high likelihood of NMD. For gRNAs that induce exon skipping by disrupting splice sites, the position of the targeted exon is provided (Figure 2B).

**Figure 2.**
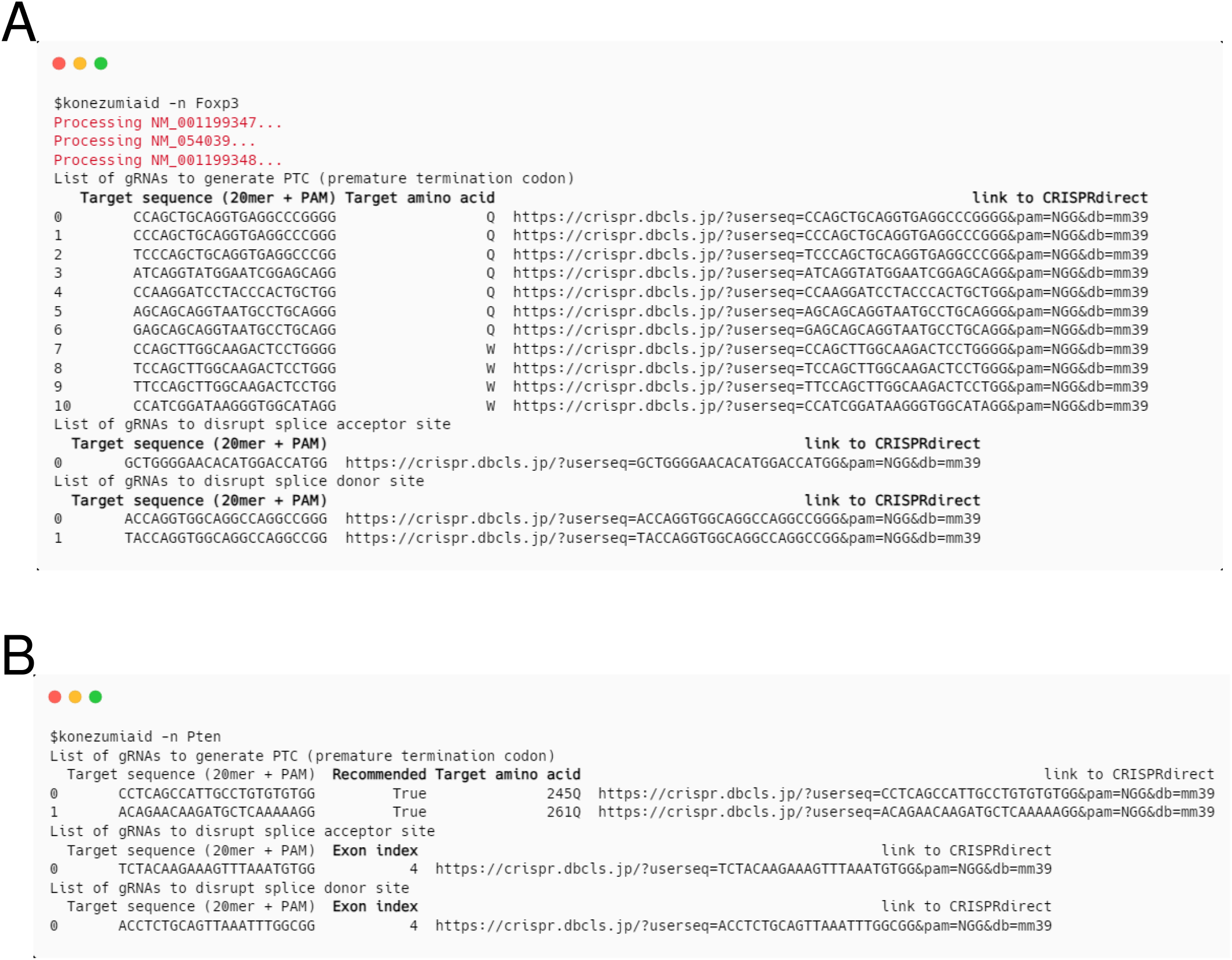
Output of KOnezumi-AID: (A) Output of KOnezumi-AID for a multi-isoform gene. The output contains gRNAs that induce PTC and gRNAs that disrupt acceptor or donor consensus nucleotides for genes with multiple isoforms. The red highlights indicate that multiple isoforms are being searched simultaneously. The bold highlights emphasize the output columns. (B) Output of KOnezumi-AID for a single-isoform gene. The output contains gRNAs that induce PTC and gRNAs that disrupt acceptor or donor consensus nucleotides for a gene with a single isoform. The bold highlights emphasize the columns that are specifically displayed when searching for single-isoform genes.

### 2.2. Gene Targetability Assessment using KOnezumi-AID

The development of KOnezumi-AID enabled us to comprehensively assess whether gRNAs can be designed to meet the KO strategy using Target-AID. A total of the 20,731 mouse genes, corresponding to protein-coding genes included in the filtered refFlat (see details in Materials and Methods), were searchable by KOnezumi-AID, of which 17,906 genes were identified as targetable genes with at least one gRNA designable. In detail, 16,818 genes had gRNAs capable of inducing a PTC, while 12,474 genes can be targeted with gRNAs that induce exon skipping by disrupting splice sites (Figure 3A). When examining the distribution of gRNA counts per gene, the largest group consisted of genes for which gRNA design was not possible. As the number of gRNAs increased, the number of targetable genes showed a decreasing trend. yet, when calculating the total number of genes with 10 or more successfully designed gRNAs, this group accounted for the largest proportion (Figure 3B). Furthermore, an analysis of exon numbers showed that out of 2,037 single-exon genes, 1,213 were targetable with at least one gRNA. This suggests that the KO strategy we defined for single-exon genes is effective. Similarly, the high targetability observed in genes with 20 or more exons (2,483 out of 2,516) highlights the approach’s versatility across different gene structures (Figure 3C). Our analysis also revealed characteristics of non-targetable genes. These genes tend to have shorter CDS lengths and fewer exons (Figure S1A). By summing the numbers for genes with up to 5 exons, this group accounted for 61.9% of the non-targetable genes (Figure S1B). These results suggest that genes with shorter CDS lengths and fewer exons are more likely to be non-targetable. Overall, these results confirm the effectiveness and flexibility of KOnezumi-AID for targeting a wide range of genes, including both single-exon and multi-exon genes.

**Figure 3.**
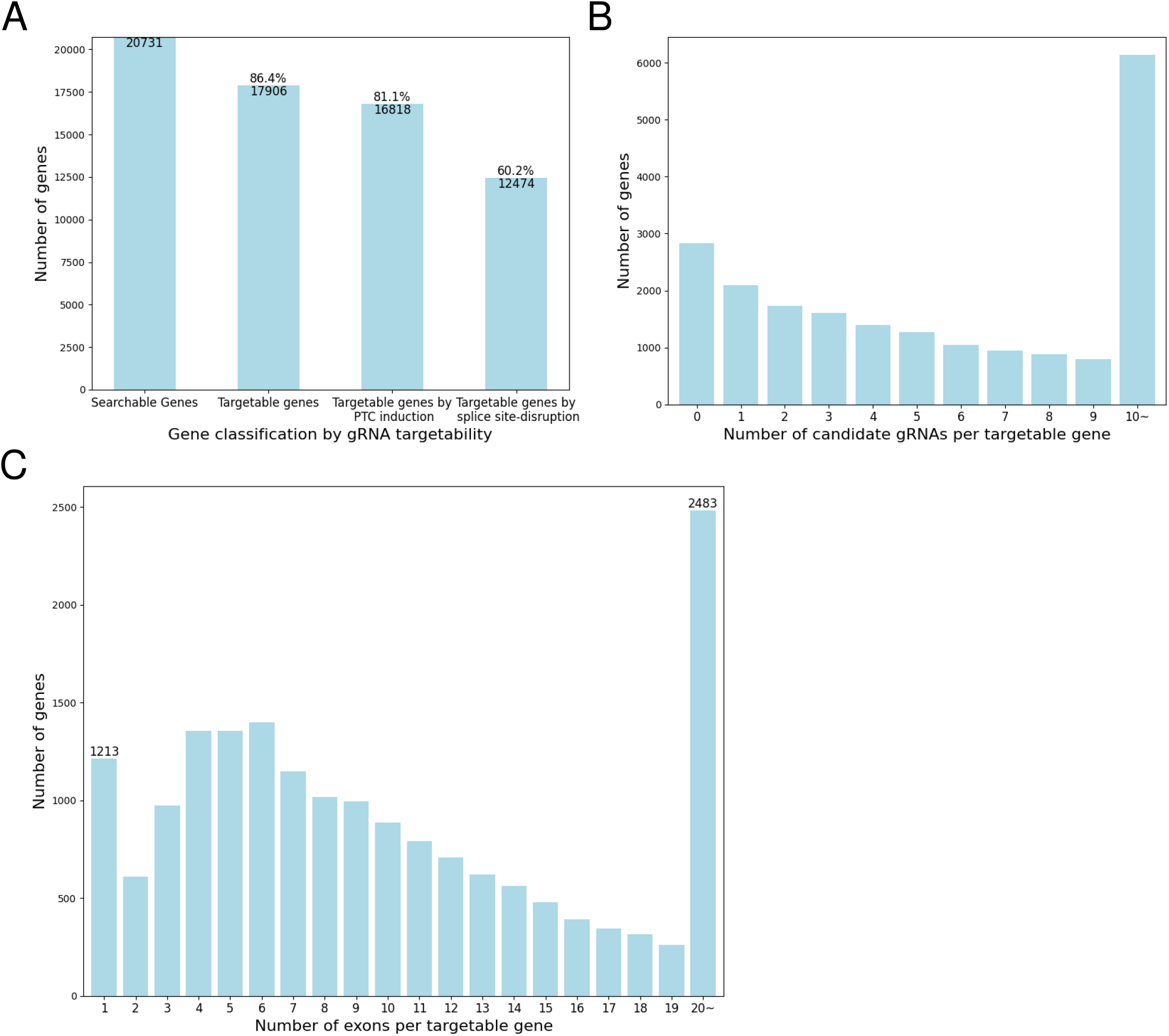
Analysis of Candidate gRNA-Associated Gene Counts and Distributions: (A) Number of genes by gRNA targetability in the mouse genome. The total number of analyzed genes (Searchable Genes), the number of genes with at least one candidate gRNA (Targetable genes), the number of genes with candidate gRNAs that induce PTC (Targetable genes by PTC induction), and the number of genes with candidate gRNAs disrupting splice sites (Targetable genes by splice site-disruption) are displayed. (B) Number of genes by number of candidate gRNAs in the mouse genome. The distribution of genes based on the total number of candidate gRNAs, including those that induce PTC and those that disrupt splice sites, is displayed. For visualization, counts are capped at 10 if the number of gRNAs per gene exceeds 10. (C) Number of genes by number of exons in the mouse genome. The distribution of exon numbers for genes that can have gRNAs designed is displayed. For visualization, counts are capped at 20 for genes with more than 20 exons.

### 2.3. Application of KOnezumi-AID to Human genome Data

The gRNA design strategy of KOnezumi-AID is not limited to mouse and can also be applied to across various species, including animals and plants, as long as their genomes have annotated exons. Therefore, to verify the robustness of the KOnezumi-AID workflow, we applied the same analysis to human genes as we did for the entire set of searchable mouse genes. In particular, we evaluated targetability, the distribution of gRNAs per gene, and the number of exons in genes where gRNAs were designed. The results revealed that KOnezumi-AID can also be used to design gRNAs for KO using Target-AID. A total of the 19,073 human genes analyzed, 16,307 were identified as targetable with at least one designable gRNA using Target-AID. Specifically, 15,112 genes had gRNAs capable of inducing PTCs, while 11,448 genes were targetable for exon skipping by disrupting splice sites (Figure 4A). The characteristics of genes with designed gRNAs in humans mirrored those observed in mice (Figure 4B, C), indicating that the design logic of KOnezumi-AID functions properly in human genome analysis as well. These findings demonstrate that the KOnezumi-AID workflow is applicable to human genome data, expanding its utility to other species as well.

**Figure 4.**
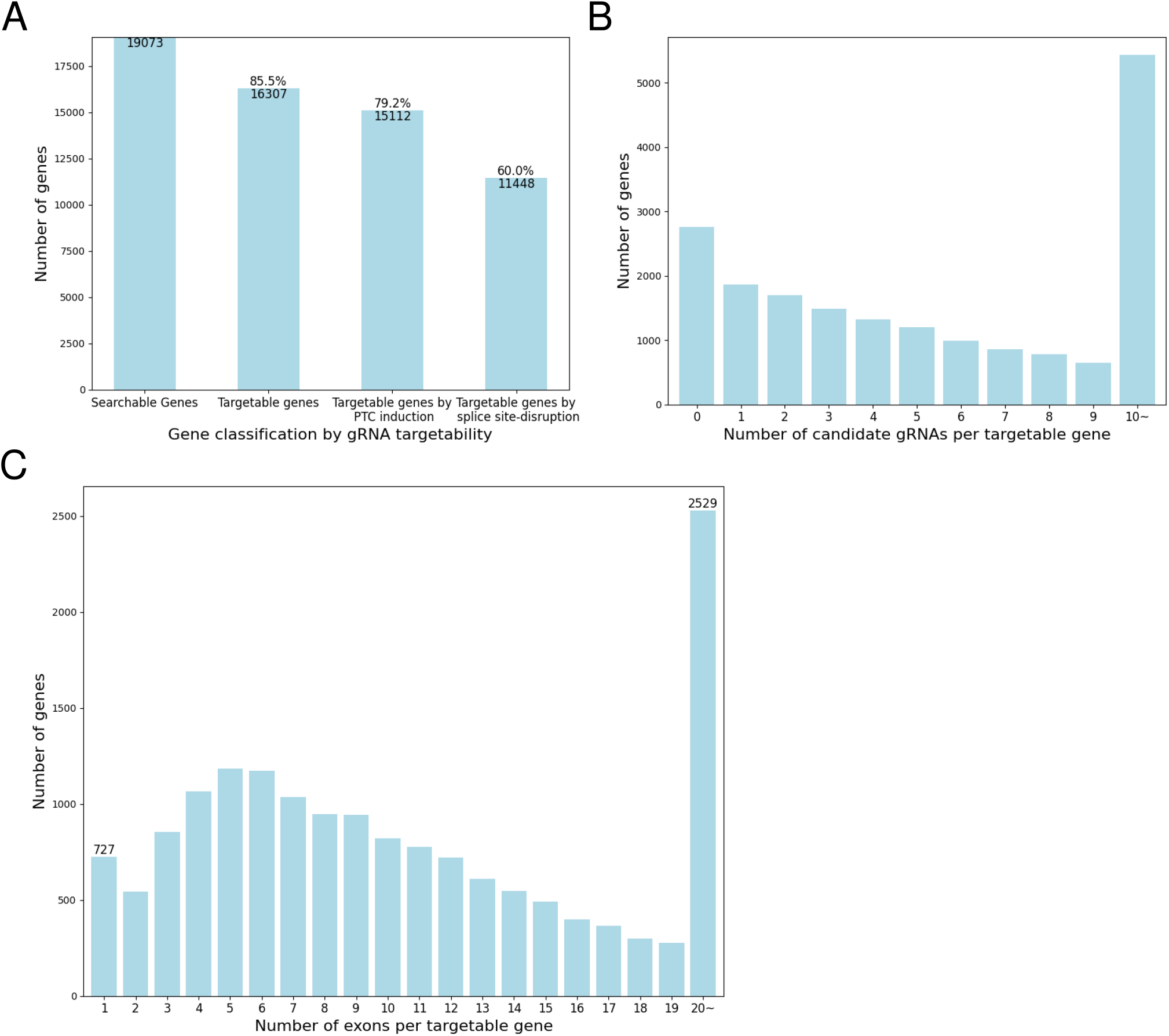
Application of KOnezumi-AID for Human Gene Data: (A) Number of genes by gRNA targetability in the human genome. The total number of analyzed genes (Searchable Genes), the number of genes with at least one candidate gRNA (Targetable genes), the number of genes with candidate gRNAs that induce PTC (Targetable genes by PTC induction), and the number of genes with candidate gRNAs that disrupt splice sites (Targetable genes by splice site-disruption) are displayed. (B) Number of genes by number of candidate gRNAs in the human genome. The distribution of genes based on the total number of candidate gRNAs, including those that induce PTC and those that disrupt splice sites, is displayed. For visualization, counts are capped at 10 if the number of gRNAs per gene exceeds 10. (C) Number of genes by number of exons in the human genome. The distribution of exon numbers for genes that can have gRNAs designed is displayed. For visualization, counts are capped at 20 for genes with more than 20 exons.

### 2.4. Batch Processing Capability of KOnezumi-AID

KOnezumi-AID streamlines the gRNA design process, that is suitable for large-scale CRISPR screening projects where multiple genes are simultaneously targeted. CRISPR screening is a method for analyzing gene functions by targeting multiple genes under specific biological conditions [27]. To facilitate such large-scale projects, we implemented a batch processing feature in KOnezumi-AID. The user prepares a CSV or Excel file devoid of any headers, containing the gene symbols or RefSeq IDs of interest. Upon executing KOnezumi-AID with the file, users can retrieve search results for multiple genes concurrently (Figure 5). This batch processing functionality allows users to efficiently design gRNAs for numerous genes, enabling quick and accurate gRNA design for large-scale genome editing projects.

**Figure 5.**
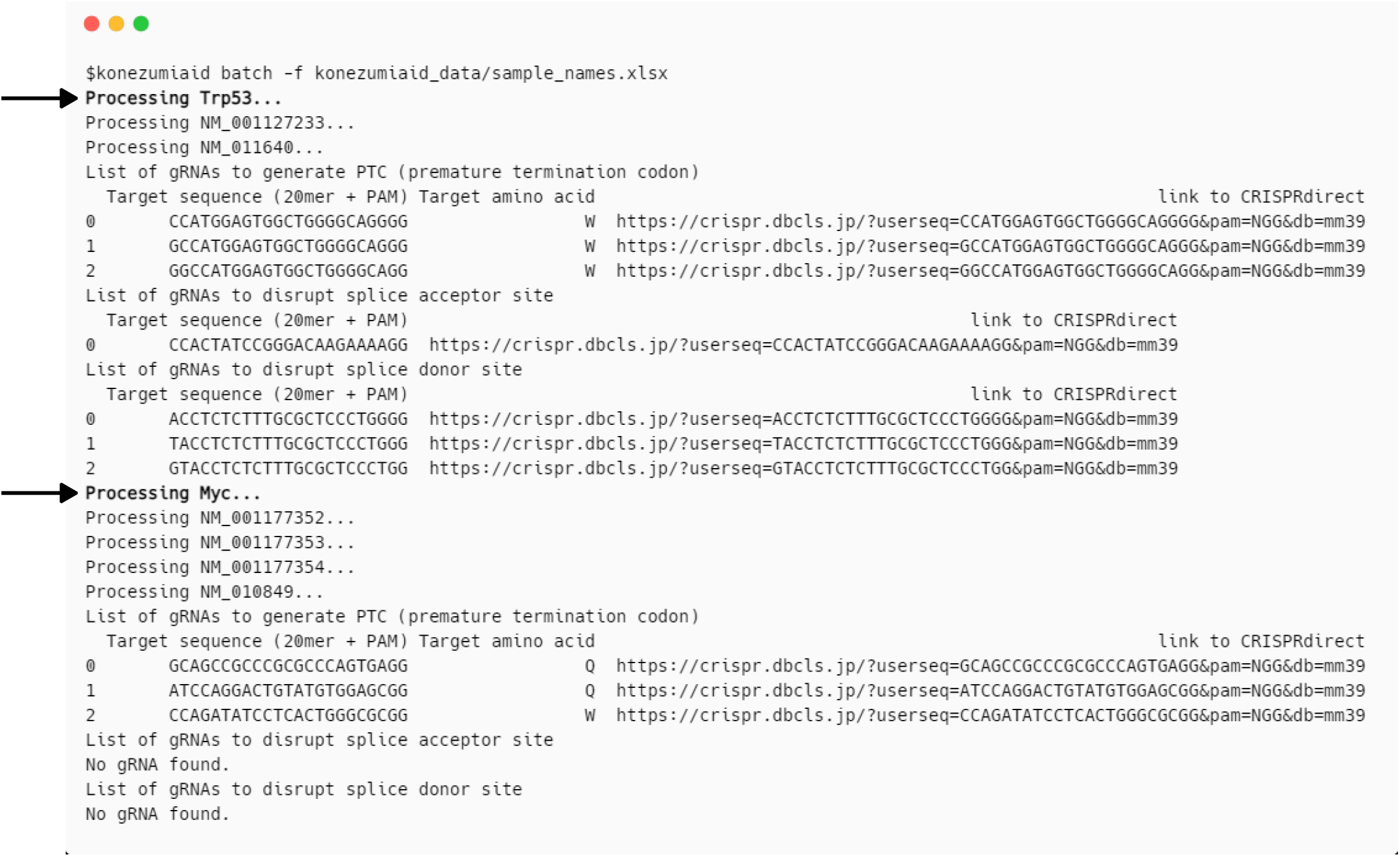
Batch Processing: Output of KOnezumi-AID for batch processing. The results of batch searches using an Excel file containing the gene names Trp53 and Myc are displayed. The bold highlights and arrows indicate that two genes are being searched sequentially.

## 3. Discussion

Base editing is a recent genome-editing technique that combines CRISPR system components with additional enzymes to introduce precise point mutations into DNA or RNA, avoiding the need for DSBs. Among these methods, Target-AID has been recognized for its lower risk of off-target RNA editing [10], making the simultaneous editing of multiple genes a promising approach.

In this study, we developed KOnezumi-AID, an automated tool for designing optimal gRNAs to induce KOs using Target-AID. Unlike other gRNA design tools, such as BE-Designer [28], which provides all possible base editor target sequences in a given input sequence, and BE-SCAN [29], which simplifies selection of effective BE tools for inducing cancer-associated SNVs, rather than KOs, KOnezumi-AID allows users to design gRNAs for KOs simply by inputting the target gene symbol or RefSeq ID. Moreover, KOnezumi-AID can target 86.4% of genes in mice and 85.5% of genes in humans, respectively. The ease of use of KOnezumi-AID, combined with the high targeting potential of Target-AID, suggests that KOnezumi-AID is a key tool for efficiently and reproducibly designing gRNAs.

This study has several limitations. The most significant limitation is the user interface. Compared to the original KOnezumi tool, which offers a user-friendly web application [30], KOnezumi-AID is currently a command line tool, which can only be used by those familiar with terminal operations. Thus, the implementation of a graphical user interface for KOnezumi-AID is an urgent task. Besides, since KOnezumi-AID currently supports only Target-AID, leaving 14% of genes without the experiment-ready gRNA designs. These genes that cannot be targeted by Target-AID could be edited by using other base editors or by using Target-AID-NG, which recognizes NG PAM sequences instead of NGG [31], potentially enabling gRNA design for all genes. Lastly, as the determinants of on-target and off-target effects in Target-AID are still unclear, further research is needed to facilitate more effective gRNA design.

In the future, integrating base editors beyond Target-AID into the KOnezumi-AID, as well as expanding its applicability to a wider range of species, will enhance versatility. Notably, the use of base editing for generating disease models through precise single-nucleotide mutations holds significant potential for advancing research into disease mechanisms. In this context, KOnezumi-AID will serve as a powerful tool to facilitate these experiments, particularly in the study of single-nucleotide variants related to human diseases.

## 4. Materials and Methods

### 4.1. Datasets

All genomic data used in this study were downloaded from UCSC Genome Browser [16]. The assembly versions of reference genome sequence and refFlat genome annotation for the mouse and human were GRCm39/mm39 (GCA_000001635.9) and GRCh38/hg38 (GCA_000001405.29), respectively. KOnezumi-AID defines non-protein-coding transcripts as those with a RefSeq ID starting with ’NR’, duplicated transcripts with the multiple same NM number for a single gene symbol, and the alternative assemblies as chromosomes including ’_alt’, ’_random’, ’_fix’, and ’_Un’ suffix.

### 4.2. Targetable splice sites

KOnezumi-AID defines splice sites according to conventional GT–AG rule, which are flanked by GT bases at the 5’ end and AG bases at the 3’ end [32]. KOnezumi-AID searches for gRNAs that include the G of the splice site, which can be editable by Target-AID on the reverse strand (G-to-A conversion), within the target window PAM (CCN) - 17 to -19 nt.

### 4.3. Statistical analysis

A two-sided Welch’s t-test was performed to compare the CDS lengths between KOnezumi-AID targetable and non-targetable genes.

### 4.4. KOnezumi-AID Download and Code Availability

The operation of KOnezumi-AID was validated on a WSL2 with Ubuntu and on macOS, using Python versions 3.9 to 3.11. The version of KOnezumi-AID used in this study was 0.3.1, which is available at http://dx.doi.org/10.5281/zenodo.12792127. KOnezumi-AID is available as an open-source software tool under the MIT license at https://github.com/aki2274/KOnezumi-AID. PyPI package is available at https://pypi.org/project/KOnezumiAID/.

## Supplementary Materials

The following supporting information can be downloaded from URL

**Supplementary Figure 1.**
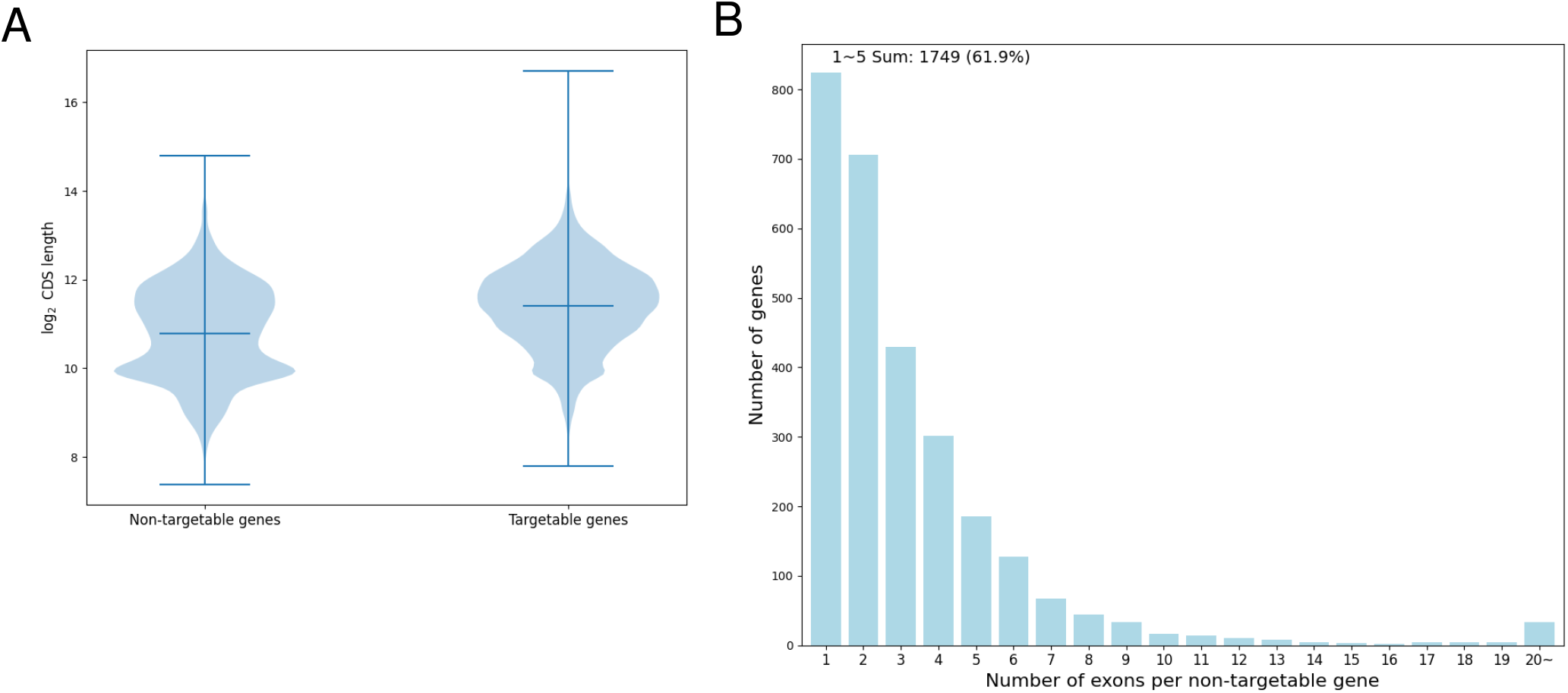
Non-targetable gene analysis: (A) This violin plot shows the distribution of log2 CDS lengths for genes with at least one candidate gRNA (targetable genes) and genes with no candidate gRNA (non-targetable genes). The solid lines in the middle of each violin represent the mean CDS length for each group, while the top and bottom edges represent the range of the data. (B) The number of exons per non-targetable gene. The distribution of the number of exons for genes with no candidate gRNA is shown. The number of exons is capped at 20 for visualization purposes. Genes with exon counts between 1 and 5 are grouped, with their total sum and percentage relative to the number of non-targetable genes displayed.

## Author Contributions

Conceptualization, S.M. and A.K.; methodology, T.T., K.M, and A.K; software, T.T and A.K.; validation, T.T.; formal analysis, T.T.; investigation, T.T., K.M and A.K; resources, T.T and A.K.; data curation, T.T and A.K.; writing—original draft preparation, T.T.; writing—review and editing, K.M,

S.M and A.K.; visualization, T.T.; supervision, S.M.; project administration, S.M and A.K.; funding acquisition, S.M. All authors have read and agreed to the published version of the manuscript.

## Data Availability Statement

The data supporting the study findings are available upon request from the corresponding authors.

## Acknowledgments

We thank Prof. Satoru Takahashi for his valuable insights and discussions regarding the project.

